# Conserved abilities of individual recognition and genetically modulated social responses in young chicks (*Gallus gallus*)

**DOI:** 10.1101/743765

**Authors:** Elisabetta Versace, Morgana Ragusa, Virginia Pallante

**Author notes:** Corresponding author, School of Biological and Chemical Sciences, Department of Biological and Experimental Psychology, Queen Mary University of London, Mile End Road 327, E1 4NS London (UK).

## Abstract

The ability to recognise familiar individuals and the motivation to stay in contact with conspecifics are important to establish social relationships from the beginning of life. To understand the genetic basis of early social behaviour, we studied the different responses to familiar/unfamiliar individuals and social reinstatement in 4-day-old domestic chicks (*Gallus gallus*) in three genetically isolated breeds: Padovana, Polverara and Robusta. All breeds showed a similar ability to discriminate between familiar and unfamiliar individuals, staying closer to familiar individuals. Social reinstatement motivation measured as the average distance between subjects, latency to the first step and exploration of the arena (a proxy for the lack of fear), differed between breeds. More socially motivated chicks that stayed in closer proximity, were also less fearful and explored the environment more extensively. These results suggest that modulation of social behaviour shows larger genetic variability than the ability to recognise social partners, which appears to be an adaptive ability widespread at the species level even for very young animals.

## 1. Introduction

In many species, the ability to recognise familiar individuals is important to establish social relationships from the first moments of life. In domestic chicks (*Gallus gallus*) and other precocial animals that move around soon after birth, the mechanism of imprinting enables to quickly learn the features of familiar individuals such as the mother and the siblings (Bateson, 1966; Bolhuis, 1991; McCabe, 2013; Vallortigara & Versace, 2018). After imprinting, hatchlings exhibit strong affiliative responses towards the imprinting stimulus while avoiding unfamiliar stimuli, a behaviour that requires the ability to recognise familiar individuals. Moreover, separation from the imprinting objects elicits attempts to signal and join the social partners (Jones & Merry, 1988; B. R. Jones & Williams, 1992; Suarez & Gallup, 1983; Zajonc, Wilson, & Rajecki, 1975), as well as adrenocortical activation in domestic chicks (Jones & Williams, 1992). Imprinting is not limited to general species recognition, as initially hypothesised by Lorenz, but enables to discriminate between particular individuals. This is shown by the different affiliative and aggressive responses to familiar and stranger social partners of the same chicken breed (Rajecki, Ivins, & Rein, 1976; Väisänen & Jensen, 2004; Vallortigara, 1992; Vallortigara, Cozzutti, Tommasi, & Rogers, 2001; Vallortigara & Andrew, 1994; Zajonc et al., 1975 see also Schweitzer, Poindron, & Arnould, 2009 for quails). Moreover, Johnson and Horn (1987) have shown that the ability of chicks to imprint on a specific hen depends on the intermediate and medial mesopallium (IMM) and Town (2011) has shown that this area responds differently to familiar and unfamiliar conspecifics.

For wild hatchlings, the ability to imprint and recognise familiar individuals is important to identify the care giver and maintain flock/group cohesion. However, recognition of familiar individuals has been documented also in hatchlings of solitary species, such as land tortoises of different species (Versace, Damini, Caffini, & Stancher, 2018). Filial imprinting is particularly suitable to investigate in the laboratory since imprinting can be established also for artificial objects that are more easily manipulated in appearance than living animals. Thanks to these features, imprinting has become a model system for memory, learning and social behaviour (Bateson, 1966; Bolhuis, 1991; Horn, 1985, 2004; McCabe, 2013, 2019). In the laboratory, imprinting is studied using well controlled artificial stimuli such as balls, cubes, cylinders or two-dimensional stimuli presented on cardboards or computer screens (Rosa-Salva et al., 2018; Versace, Schill, Nencini, & Vallortigara, 2016; Versace, Spierings, Caffini, ten Cate, & Vallortigara, 2017; Wood & Wood, 2015). These experiments have shown that chicks discriminate subtle differences of the imprinting objects, such as rotation of the features located inside the imprinting object (Vallortigara & Andrew, 1991), the configuration of items that compose the imprinting stimulus (Rosa-Salva et al., 2018) and even the underlying structure of the stimuli independent of their physical appearance (Martinho & Kacelnik, 2016; Versace, Regolin, & Vallortigara, 2006; Versace, Spierings, et al., 2017). Less is known on the genetic differences in the early ability of chicks to imprint on living objects and to discriminate between familiar and unfamiliar individuals. Väisänen and Jensen (2004) have explored the differences in responses to familiar and unfamiliar social stimuli in red jungle fowls (a breed considered to be close to the ancestral undomesticated chicken (Miao et al., 2013) and White Leghorns (a modern breed selected for laying eggs), using 3-4 week old animals. This study showed greater affiliative responses in White Leghorn compared to red jungle fowls but did not clarify the onset of these differences.

Here we explore the genetic differences of chicks in responding to familiar/unfamiliar individuals in 4-day old chicks of three different breeds of domestic fowl: the Padovana, Polverara and Robusta breed. Chicks originated in the same farm within the conservation programme Co.Va, that kept these local breeds in genetic isolation for more than 20 years (De Marchi, Cassandro, Targhetta, Baruchello, & Notter, 2005). This particular arrangement reduced the environmental differences while ensuring low admixture between breeds (Zanetti, De Marchi, Dalvit, & Cassandro, 2010). We previously investigated the predisposed visual preferences of these breeds to approach a stuffed hen vs. a scrambled version of it in visually naïve chicks (Versace, Fracasso, Baldan, Dalle Zotte, & Vallortigara, 2017). Predisposed responses precede imprinting for they are exhibited before any prior visual experience has occurred and do not depend on experience (Di Giorgio et al., 2017). When given the choice between a stuffed hen and a scrambled version of it, we observed that all three breeds initially preferred to orient towards the stuffed hen, which is the predisposed stimulus that several breeds preferentially approach (see Egorova & Anokhin, 2003; Johnson & Horn, 1988; Mayer, Rosa-Salva, Lorenzi, & Vallortigara, 2016; Versace, Fracasso, Baldan, Dalle Zotte, & Vallortigara, 2017). Interestingly, we observed a difference between breeds as early as the first 10 minutes of visual experience: while the Polverara breed showed a steady attachment for the stuffed hen throughout the test, the Robusta and Padovana breeds were attracted by the alternative stimulus. This suggests that behavioural strategies that drive early attachment and orientation have a genetic basis, and that genetic differences are apparent in the first minutes of visual experience. It is not clear, though, whether differences in the ability to recognise familiar individuals and social motivation are present between these breeds. The peculiar skull and brain anatomy of the Padovana breed is a further reason of interest for these breeds. Both the Padovana and Polverara breed have a striking crest on the head but only in the Padovana breed the crest covers a perforated skull with an associated enlargement of the brain (Verdiglione & Rizzi, 2018). The behavioural and cognitive implications of this striking anatomy remain elusive, although Darwin (1868) hypothesised potential deficits in a closely related breed, the white crested Polish. Historical documents suggest that the Padovana breed, whose traces go back to Roman times (Brothwelp, 1979), was introduced in Italy from Poland more than seven centuries ago (De Marchi et al., 2005), originating from the white crested Polish. In the White crested Polish/Padovana breed the endocranium is enlarged and the brain has dramatically expanded to fill this gap (Frahm & Rehkamper, 1998). It is not clear which behavioural consequences this peculiar brain organisation has produced, and whether the abilities of individual recognition and affiliative responses for this breed differ from those of other chickens.

To investigate the genetic variability in early individual recognition and in social motivation we tested the three breeds in an open field test. In this setting, chicks are located in a novel empty arena larger than their home cage and their spontaneous behaviour is observed in the presence of familiar or stranger conspecifics. The ability of chicks to recognise familiar/stranger individuals can be inferred looking at whether the distance kept between familiar animals is different. Due to the process of filial imprinting, familiar chicks are expected to stay closer than stranger chicks, as previously documented (see for instance Vallortigara, 1992; Zajonc et al., 1975). Based on their antipredatory and affiliative behaviour, it has been suggested that animals located in an open field with other conspecifics experience both the fear of being in an open environment – which has the effect of reducing activity and exploration –, and social reinstatement, namely the motivation to reach the group and remain in contact with conspecifics (Suarez & Gallup, 1983; Vallortigara, 1992; Vallortigara, Cailotto, & Zanforlin, 1990). In a range of species, greater latency of movement/tonic immobility indicates antipredatory responses, while shorter distance between individuals indicates stronger social/reinstatement motivation (Jones, Mills, & Faure, 1996; Versace, Caffini, Werkhoven, & Bivort, 2019).

## 2. Methods

### 2.1 Breeds and conservation scheme

We studied three genetically isolated breeds of domestic fowl (*Gallus gallus*) raised in the same farm (Istituto Istruzione Superiore Agraria “Duca degli Abruzzi”, Padova, Italy): Padovana, Polverara and Robusta maculata. These breeds entered the Co.Va conservation project more than 20 years before this project (De Marchi et al., 2005). The breeding and conservation scheme included no gene flow between breeds and were aimed at increasing the number of pure breed animals while maintaining genetic variability within the breed. We included individuals from gold, silver and buff Padovana and white and black Polverara, because previous studies showed high homogeneity within these breeds (Zanetti, De Marchi, Abbadi, & Cassandro, 2011; Zanetti et al., 2010). The Robusta maculata breed was developed in 1965 from crosses between Tawny Orpingtons and White Americans. Zanetti et al. (2010) documented genetic isolation (low level of admixture) between these breeds and a closer phylogenetic relationship between Padovana and Polverara, which are also more similar at phenotypic level compared to Robusta. More details are provided in Versace et al. (2017).

### 2.2 Subjects and rearing conditions

We tested 221 pairs of chicks: 94 pairs of the Padovana breed (PD), 88 pairs of the Polverara breed (PL), 39 pairs of the Robusta breed (RB). During the test, 156 pairs moved: 58 Padovana, 60 Polverara and 38 Robusta. Eggs were obtained from the Agricultural High School “Duca degli Abruzzi” (Padova, Italy), which is pursuing the Co.Va conservation programme for the maintenance of local biodiversity described above (De Marchi et al., 2005). We incubated and hatched eggs in darkness at 37.7 °C. During incubation, humidity was kept at 40% and then increased to 60% during the last three days of incubation. Chicks hatched in groups of the same breed and were then housed in pairs of the same breed within 24 hours from hatching without any visual exposure to conspecifics prior to housing. Chicks were maintained in standard rearing conditions (temperature 28°C, 14:10 day:night cycle) for three days in 28×38×32 cm cages, and tested at an age of 4 days after hatching. After the experiments, animals were donated to local farmers.

### 2.3 Apparatus

The test apparatus used for the test was a black square area (40×40×36 cm), illuminated with an incandescent lamp (100 W) located 1 m above the centre of the arena. A camera recorded the experimental scene from above during the test.

### 2.4 Test procedure

The experiment consisted of two phases: a familiarization phase and a test phase (see Figure 1). The familiarization phase started soon after hatching (day post hatching 0) and lasted 3 days. During this phase a pair of chicks of the same breed was housed in the same cage. At test, each subject was visually isolated for 10-15 minutes in an opaque box (14.5 cm height, 8.5 cm large, 11 cm width) located inside its cage and then transferred to the testing room together with another experimental subject. During the test we observed two chicks previously housed together (Familiar condition) or previously housed with a different animal (Stranger condition). At the beginning of the test, experimental chicks were placed simultaneously in two opposite corners of the arena and left free to move for 5 minutes.

**Figure 1.**
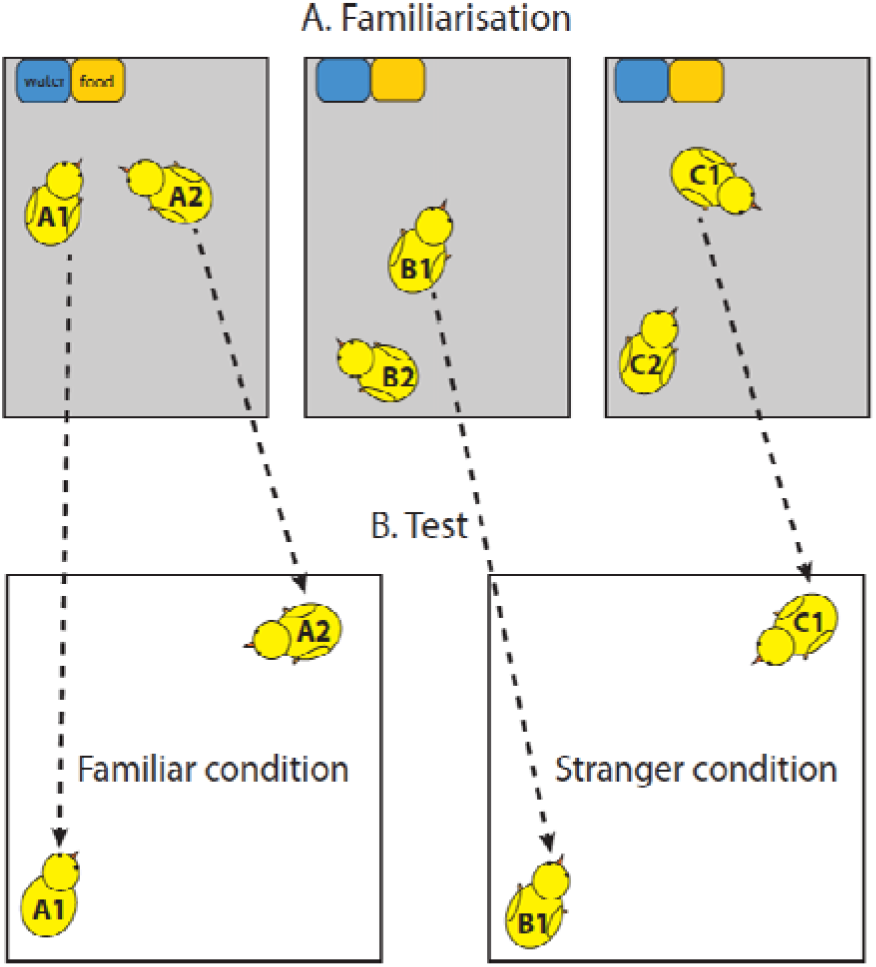
(A) Chicks were initially housed with another chick of the same breed for three days (familiarisation). (B) At test, chicks were located in a new arena either with the familiar companion (Familiar condition) or with a novel chick (Stranger condition). For each pair we recorded the latency of the first step, the distance between the centroid of the chicks and overall distance run.

### 2.5 Data analysis

We initially established whether chicks in the pair moved or remained still throughout the test and counted how many pairs moved/did not move during the test. We used a Chi-square test to check whether more pairs moved in the Stranger or in the Familiar condition.

For all pairs that moved, we used an Anova to analyse the average distance (cm) between the centroid of subjects throughout the test, the overall distance moved by the pair (cm) and the average latency of the first step (s) using Condition (familiar, stranger), Breed (Padovana, Polverara, Robusta) and their interaction as independent variables. Exploratory analyses showed that a log-transformation of the distance moved and latency normalised the residuals of these variables, hence we used log-transformed values for these analyses. We ran post-hoc t-tests to explore significant differences. We also analysed the relation between latency, distance run and distance between subjects using the Spearman rank order correlation test and fitting linear and polynomial models until finding the minimal adequate model. Significance level was set to p<0.05. Statistical analyses were performed with the R software (version 3.5.2).

## 3 Results

### 3.1 Pairs that moved in the Familiar and Stranger condition

There was no significant difference in the frequency of pairs that moved/did not move between the Familiar and Stranger condition (X-squared=0.834, df=1, p=0.361) or Breed (X-squared=0.989, df=2, p=0.610).

### 3.2 Distance between individuals

Looking at the log-transformed distance between subjects we found a significant effect of Condition (F_1,150_=8.311, p=0.005) and Breed (F_2,150_=5.664, p=0.004) and no significant interaction (F_2,150_=0.155, p=0.857). These results are shown in Figure 2. Familiar chicks stayed closer than stranger chicks (t=−2.817, df=153.34, p=0.005). The average distance between Familiar chicks was 17.57 cm (median 10.26 cm), the average distance between Stranger chicks 20.75 cm (median 15.35 cm).

**Figure 2.**
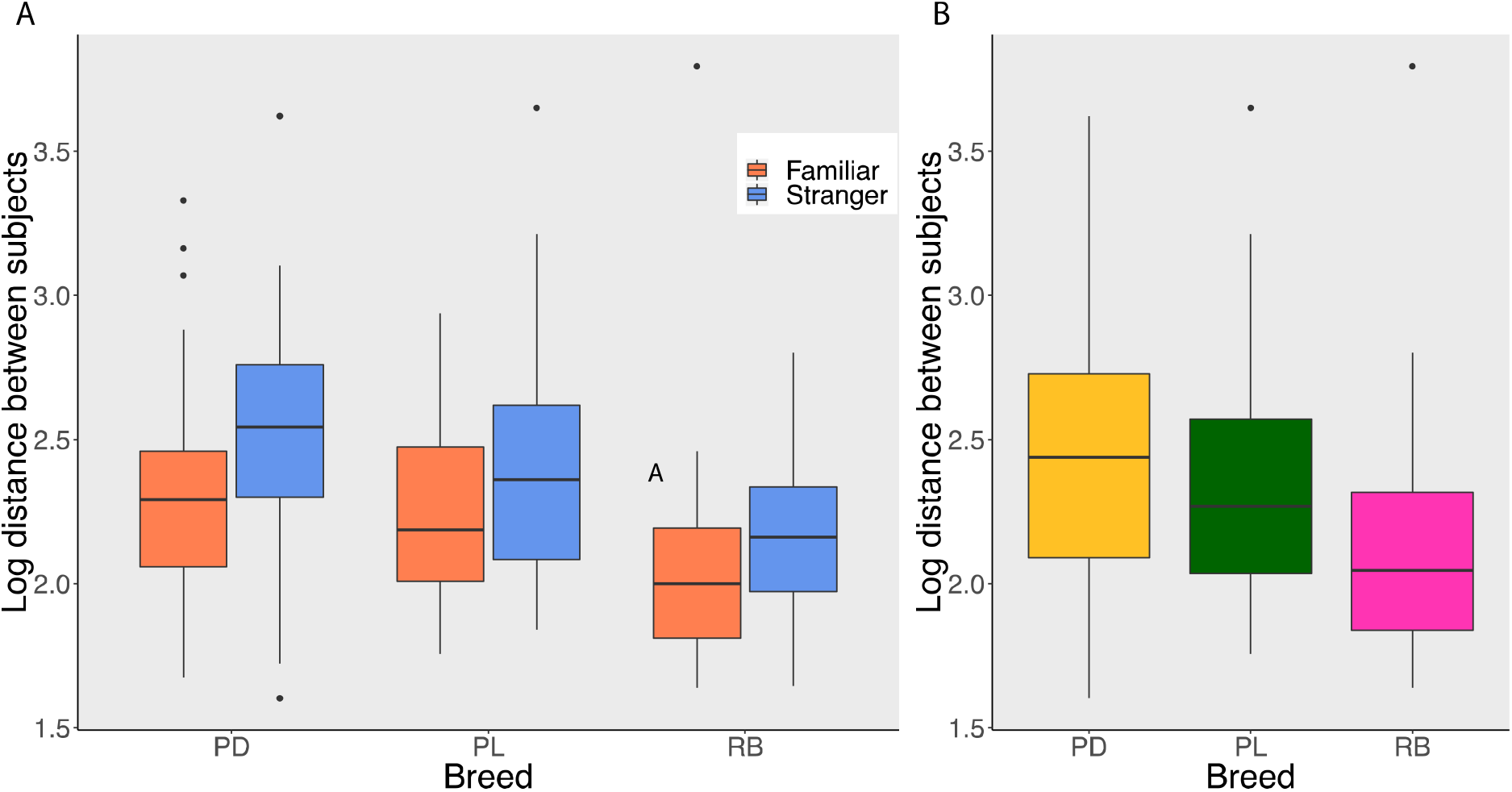
The boxplots show (A) for each condition (Familiar, Stranger) and (B) breed (PD=Padovana, PL=Polverara, RB=Robusta) the median, quartiles, maximum, minimum and outliers of the distance between subjects expressed in log centimetres.

Chicks in the RB breed stayed significantly closer than PL (t=2.536, df=76.942, p=0.013) and PD (t=3.455, df=80.728, p<0.001) chicks, whereas there was no significant difference between PL and PD chicks (t=1.139, df=115.01, p=0.257).

### 3.3 Distance run

Analysing the overall distance run in each pair, we found a significant effect of Breed (F_2,150_=6.102, p=0.003) and no significant effect of Condition (F_1,150_=0.409, p=0.523) and no significant interaction (F_2,150_=1.027, p=0.360). These results are shown in Figure 3.

**Figure 3.**
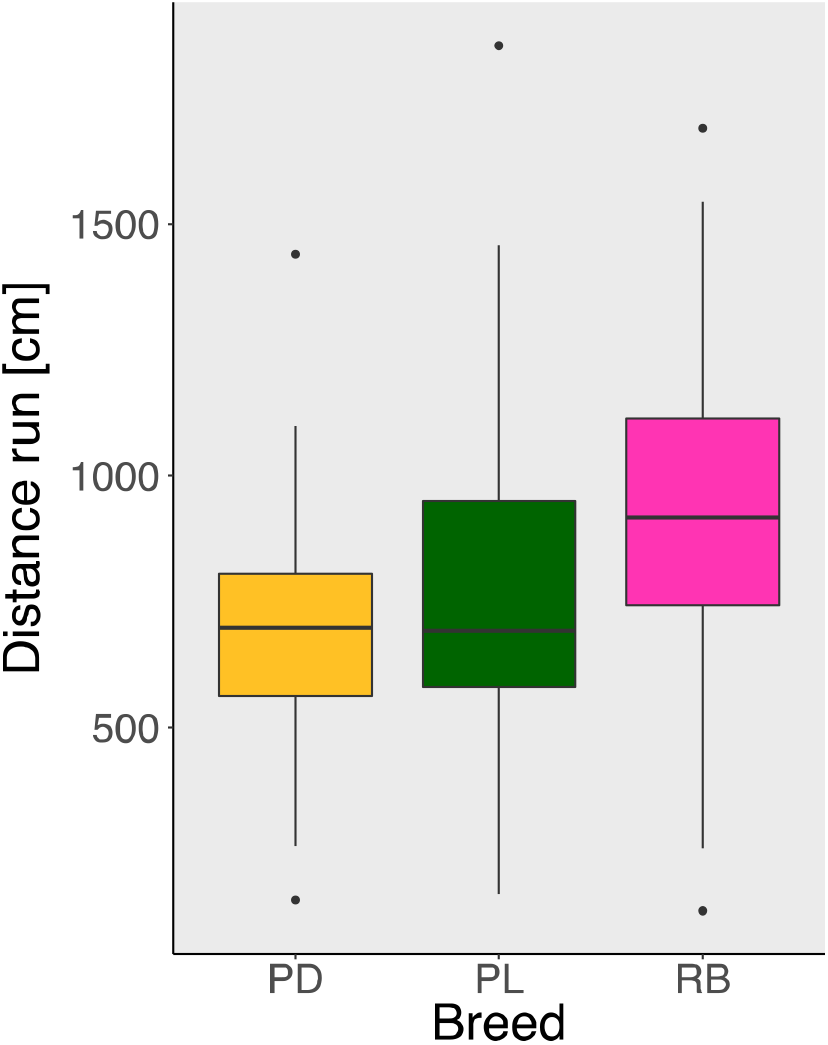
The boxplots show for each breed (PD=Padovana, PL=Polverara, RB=Robusta) the median, quartiles, maximum, minimum and outliers of the distance run expressed in centimetres.

Polverara chicks did not run significantly more than Padovana chicks (t= 1.890, df=110.13, p=0.061) although there was a trend in this direction, Robusta chicks ran significantly more than Polverara chicks (t=2.530, df=71.406, p=0.014), and significantly more than Padovana chicks (t=4.186, df=58.757, p<0.001).

### 3.4 Latency first step

Analysing the latency of the first step in each pair, we observed a significant effect of Condition (F_2,150_=4.026, p=0.047, with chicks in the Familiar condition moving sooner than chicks in the Stranger condition) and Breed (F_2,150_=11.02, p<0.001) and no significant interaction (F_2,150_=0.336, p=0.715). The mean latency for the Familiar condition was 43.29 s (median 20.25 s) and for the Stranger condition 48.19 s (median 36 s). The results are shown in Figure 4.

**Figure 4.**
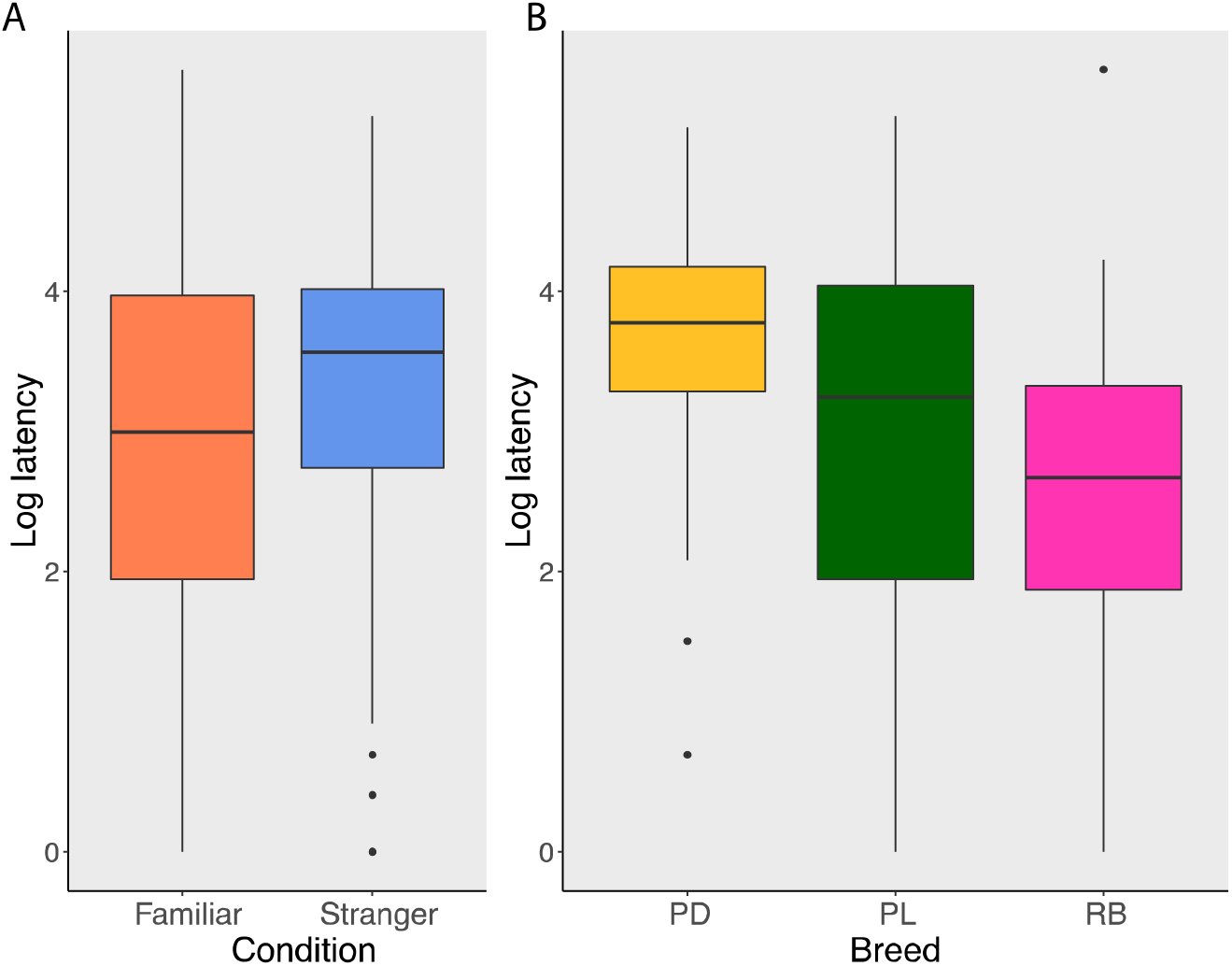
The boxplots show for each (A) condition (Familiar, Stranger) and (B) breed (PD=Padovana, PL=Polverara, RB=Robusta) the median, quartiles, maximum, minimum and outliers of the pair latency of the first step expressed in log seconds.

Polverara chicks moved significantly earlier than Padovana chicks (t=-27.937, df=78.274, p<0.001), Robusta chicks moved significantly earlier than Polverara chicks (t=-27.229, df=71.209, p<0.001), and also Robusta chicks moved significantly earlier than Padovana chicks (t=-25.87, df=42.701, p<0.001).

### 3.5 Relation between latency, distance between subjects and distance run

We observed a significant positive correlation between the latency of the first step and the average distance between subjects (S= 162850, p<0.001, rho=0.743, see Figure 5A) and a significant negative correlation between the latency of the first step and the distance run (S=899280, p<0.001, rho=-0.421, see Figure 5B).

**Figure 5.**
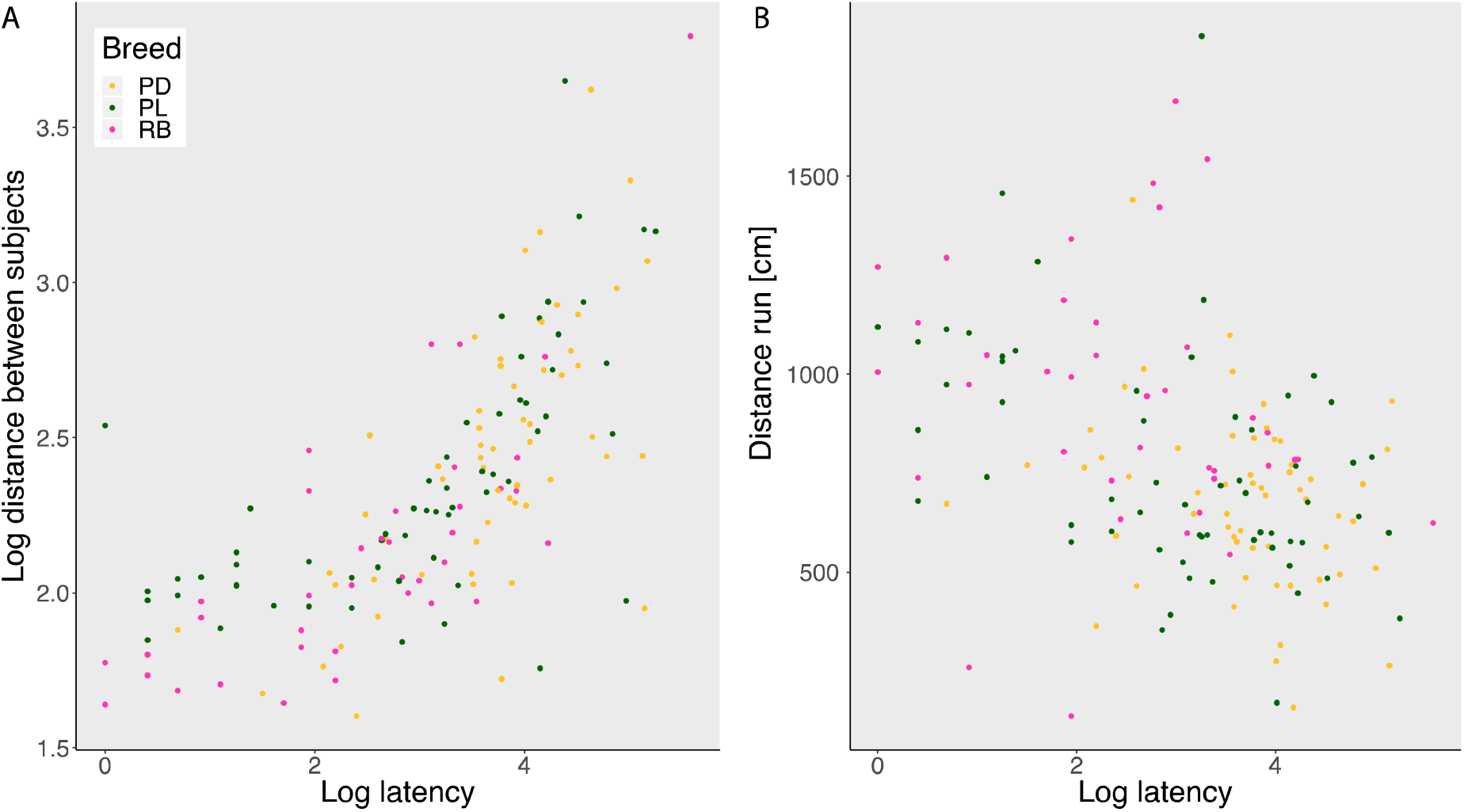
The scatterplots show the relation between (A) latency of the first step and average distance between subjects and (B) latency of the first step and distance run.

The minimal adequate model for the dependent variable Distance between subjects using as predictors Latency, Condition and Breed included a quadratic polynomial with Latency (F_2,150_=98.035, p<0.01), Condition (F_1,150_=7.13, p=0.008) and Latency x Condition (F_2,150_=4.77, p=0.001) as significant effects. The full table of coefficients is reported in Table 1.

**Table 1.** This table shows the coefficients of the minimal adequate model for the dependent variable Distance between subjects using as predictors Latency, Condition and Breed.

|  | Estimate | Std. Error | T value | Pr(> \|t\|) |
| --- | --- | --- | --- | --- |
| Intercept | 2.266 | 0.032 | 70.122 | <2e-16 |
| Latency (linear) | 3.138 | 0.370 | 8.493 | 1.83e-14 |
| Latency (quadratic) | 0.886 | 0.366 | 2.421 | 0.017 |
| Condition (Stranger) | 0.120 | 0.045 | 2.642 | 0.009 |
| Latency (linear) x Condition | 1.135 | 0.576 | 1.970 | 0.051 |
| Latency (quadratic) x Condition | 1.373 | 0.572 | 2.400 | 0.018 |
| Residual standard error: 0.2797 on 150 degrees of freedom |  |  |  |  |
| Multiple R-squared: 0.587 |  | Adjusted R-squared: 0.573 |  |  |
| F-statistic: 42.55 on 5 and 150 df, p<2.2e-16 |  |  |  |  |

The minimal adequate model for the dependent variable Distance run using as predictors Latency, Condition and Breed included Latency (F_1,151_=29.100, p<0.01) and Breed (F_2,151_=4.322, p=0.015) as significant effects. The full table of coefficients in reported in Table 2.

**Table 2.**
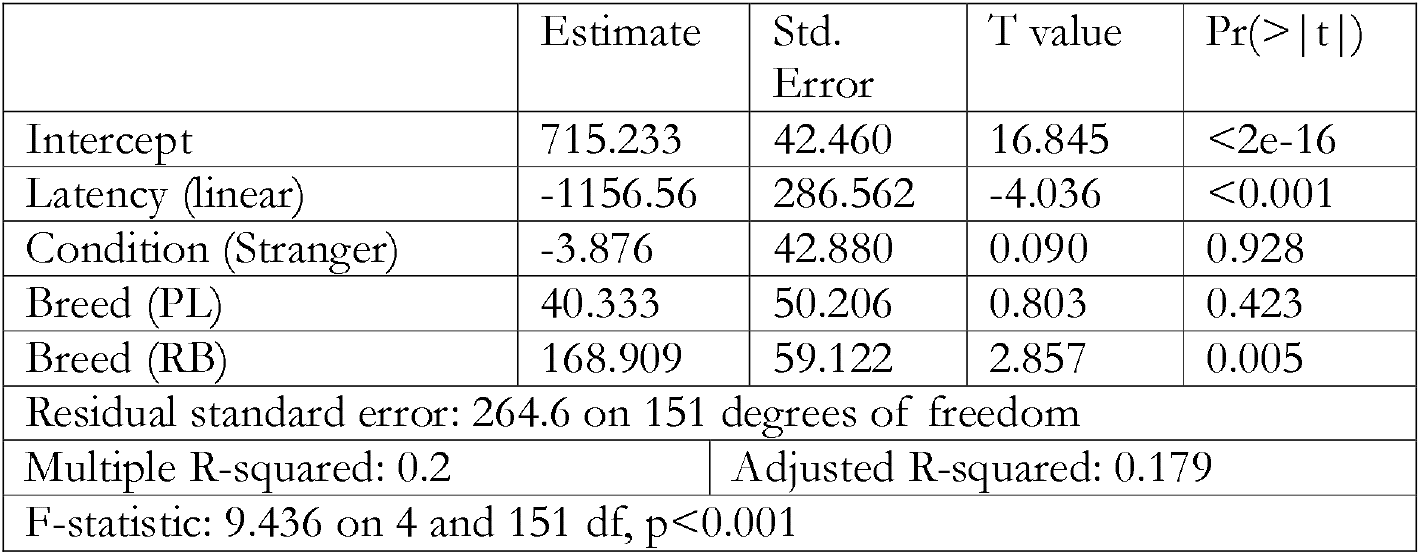
This table shows the coefficients of the minimal adequate model for the dependent variable Distance run using as predictors Latency, Condition and Breed.

|  | Estimate | Std.<br>Error | T value | Pr(> \|t\|) |
| --- | --- | --- | --- | --- |
| Intercept | 715.233 | 42.460 | 16.845 | <2e-16 |
| Latency (linear) | -1156.56 | 286.562 | -4.036 | <0.001 |
| Condition (Stranger) | -3.876 | 42.880 | 0.090 | 0.928 |
| Breed (PL) | 40.333 | 50.206 | 0.803 | 0.423 |
| Breed (RB) | 168.909 | 59.122 | 2.857 | 0.005 |
| Residual standard error: 264.6 on 151 degrees of freedom |  |  |  |  |
| Multiple R-squared: 0.2 |  | Adjusted R-squared: 0.179 |  |  |
| F-statistic: 9.436 on 4 and 151 df, p<0.001 |  |  |  |  |

## 4. Discussion

From the beginning of life, the ability to recognise familiar individuals and the motivation to stay in contact with conspecifics are important to establish social relationships. Little is known, though, on whether early social behaviours have a genetic basis that determines behavioural differences. To address this issue, we focused on domestic chicks (*Gallus gallus*). Chicks are an ideal model to investigate the first social responses not only because they are a precocial species (Versace & Vallortigara, 2015; Versace, 2017) and move around autonomously soon after birth but also because they live in flocks and have to recognise the mother and siblings to maintain social cohesion (Nicol, 2015). We previously showed that the initial predisposition to orient towards a (stuffed) hen does not vary between breeds, while exploratory responses to social stimuli differ between breeds after just 10 minutes of experience (Versace, Fracasso, Baldan, Dalle Zotte, & Vallortigara, 2017). Here, we investigated early social responses mediated by previous social experience and learning. We tested whether 4-day-old chicks have standing genetic variation for the ability to recognise conspecifics and social reinstatement by using three genetically isolated breeds: Padovana, Polverara and Robusta.

In all breeds, we observed a similar ability to discriminate between particular individuals and recognise a familiar one. As a proxy for recognition, we used the different distance kept between familiar and unfamiliar individuals. When located in a new environment with a familiar or an unfamiliar chick, all tested breeds equally discriminated between familiar and unfamiliar individuals staying significantly closer to familiar chicks. and showing shorter latency of movement with pairs of familiar individuals. Even the Padovana breed, that shows the neuroanatomical peculiarity of a perforated skull with an associated enlargement of the brain (Verdiglione & Rizzi, 2018), did not show cognitive differences in this ability. This performance of Padovana chicks suggests that the behavioural limitations noticed by Darwin (1868) in the white crested Polish – a breed ancestral to Padovana (De Marchi, Dalvit, Targhetta, & Cassandro, 2005; Verdiglione & Rizzi, 2018) – do not derive from deficits in social motivation or individual recognition (they might instead be due to difficulties in visual perception linked to the long plumage of the crest as suggested by Vallortigara, personal communication). Hence, the ability to promptly recognise familiar individuals appeared widespread at the species level in chickens. Previous studies conducted on quails (*Coturnix coturnix japonica*) (Kovach, 1990) showed a slow response to selection for low/high imprintability. In this study, imprintability was selected as low/high ability to exhibit imprinting responses for colours that were not initially preferred by young birds. The observed sluggish response to selection for imprintability observed in quails is in line with the absence of interbreed differences in the recognition of familiar individuals that we have documented in chicks.

We have previously shown that even hatchlings of non-social species such as tortoises are able to recognise familiar and unfamiliar individuals in the first days of life (Versace, Damini, Caffini, & Stancher, 2018). The early age and little experience required (see also Suarez & Gallup, 1983; Vallortigara, 1992; Zajonc et al., 1975), together with the low genetic variability of this trait, shows how crucial individual recognition is at the onset of life for a range of different species. This contrasts with other social traits that showed significant genetic variability. In fact, we observed very clear differences between breeds and conditions when looking at measures of social reinstatement and fear in an open field. We measured social reinstatement, the motivation to join the social group and keep in contact with it, looking at the average distance between subjects. Shorter distance indicates greater social reinstatement motivation (Jones et al., 1996; Schweitzer et al., 2009; Suarez & Gallup, 1983; Vallortigara et al., 1990; Vallortigara, 1992). Both the Polverara and Padovana breed, that are closely genetically related (Zanetti et al., 2010), maintained a larger distance than the Robusta breed, providing another indication of the fact that the observed differences have a genetic basis. We also assessed the fear elicited by being in a novel open field by looking at the latency to the first step and overall distance run. Since an adaptive antipredator response for chicks in an open field is freezing/tonic immobility, fear is expected to induce greater latency, and less exploration in the arena as antipredator behaviours. We observed that chicks with lower latency stayed closer and explored the arena more extensively. While fear/antipredatory responses and social reinstatement can dissociate (for instance with short distance between subjects that indicates high social reinstatement and little distance moved that indicates high fear), we observed that they were inversely correlated. This shows that pairs of chicks that approached each other explored the arena while remaining in close proximity.

Differences in social reinstatement and fear responses were apparent from the first days of life, in animals with very limited social experience. The differences in social reinstatement and fear had a clear genetic basis, with an effect of breed on latency, distance run and distance between individuals. In particular, the Robusta breed showed the shortest latency and distance between subjects as well as greater distance run during the test. These results are in line with the fact that quails respond to selection for high/low social reinstatement and fear (Formanek et al., 2008; Mills & Faure, 1991; Mills, Jones, & Michel, 1995), that older chicks of jungle fowl and White Leghorn exhibit different social reinstatement behaviour (Väisänen & Jensen, 2004), and that neonate chicks differ in exploratory responses to social stimuli (Versace, Fracasso, Baldan, Dalle Zotte, & Vallortigara, 2017).

Our findings suggest that the modulation of social behaviour shows larger genetic variability than the ability of recognizing familiar individuals. This points at the crucial adaptive role of individual recognition, an ability widespread across taxa and available from the earliest developmental stages also in tortoises (Versace, Damini, Caffini, & Stancher, 2018). Interestingly, while we have shown that chicks consistently tend to aggregate with other individuals, tortoise hatchlings – that are solitary until reaching sexual maturity – ignore familiar individuals and avoid strangers (Versace, Damini, Caffini, & Stancher, 2018). The ability to promptly recognize familiar individuals can hence sustain both affiliative and avoidance responses. The pivotal role of individual recognition is apparent in invertebrates as well. For instance, in studying aggression, Yurkovic et al. (2006) have documented the ability of fruit flies to respond differently to familiar and unfamiliar opponents. Not only familiar opponents had significantly fewer encounters but the fighting strategies depended on whether the opponent was familiar or unfamiliar: losers tested with unfamiliar winners were more aggressive than losers paired with familiar winners. Individual recognition has been documented in other insects such as paper wasps (Tibbetts, 2002; Tibbetts & Dale, 2007) and ants (D’Ettorre & Heinze, 2005). It hence appears that individual recognition has a crucial role in a broad comparative context. Using newly hatched chicks of different breeds, we have shown the ability to recognize familiar individuals and differentially respond to them is widespread in this species from the earliest stages of life. We suggest that identifying the core abilities exhibited by young animals at the beginning of life might also guide experts in artificial intelligence in understanding which are the fundamental components of general intelligence (Versace, Martinho-Truswell, Kacelnik, & Vallortigara, 2018). Further studies should clarify the role of different perceptive cues chicks in discriminating between familiar individuals.

### Animal welfare note

All experiments comply with the ASAB/ABS Guidelines for the Use of Animals in Research and with the current Italian and European Community laws for the ethical treatment of animals and the experimental procedures were approved by the Ethical Committee of University of Trento and licensed by the Italian Health Ministry (permit number 1138/2015 PR). At the end of the experiments, all chicks were donated to local farmers.

## Acknowledgements

This work was funded by the Royal Society Research Grant RGS\R1\191185.

